# Vaccination with B.1.1.7, B.1.351 and P.1 variants protects mice from challenge with wild type SARS-CoV-2

**DOI:** 10.1101/2021.08.05.455212

**Authors:** Fatima Amanat, Shirin Strohmeier, Philip Meade, Nicholas Dambrauskas, Barbara Mühlemann, Derek J. Smith, Vladimir Vigdorovich, D. Noah Sather, Lynda Coughlan, Florian Krammer

## Abstract

Vaccines against severe acute respiratory syndrome coronavirus 2 (SARS-CoV-2) have been highly efficient in protecting against coronavirus disease 2019 (COVID-19). However, the emergence of viral variants that are more transmissible and, in some cases, escape from neutralizing antibody responses has raised concerns. Here, we evaluated recombinant protein spike antigens derived from wild type SARS-CoV-2 and from variants B.1.1.7, B.1.351 and P.1 for their immunogenicity and protective effect *in vivo* against challenge with wild type SARS-CoV-2 in the mouse model. All proteins induced high neutralizing antibodies against the respective viruses but also induced high cross-neutralizing antibody responses. The decline in neutralizing titers between variants was moderate, with B.1.1.7 vaccinated animals having a maximum fold reduction of 4.8 against B.1.351 virus. P.1 induced the most cross-reactive antibody responses but was also the least immunogenic in terms of homologous neutralization titers. However, all antigens protected from challenge with wild type SARS-CoV-2 in a mouse model.

**Author Summary:** The emergence of variants of SARS-CoV-2 has led to an urgency to study whether vaccines will lead to cross-protection against these variants. Here, we demonstrate that vaccination with spike proteins of various variants leads to cross-neutralizing responses, as well as protection in a mouse model against wild type SARS-CoV-2.

## Introduction

Severe acute respiratory syndrome coronavirus 2 (SARS-CoV-2) emerged in late 2019 in Wuhan, China. Since then, the virus has caused the coronavirus disease 2019 (COVID-19) pandemic leading to approximately 4 million official deaths globally (as of July 2021). While coronaviruses usually mutate more slowly than other RNA viruses due to the proof-reading activity of their replication machinery [1], viral variants started to emerge in the summer of 2020 in humans and mink in Europe [2–4]. In late 2020, additional variants, termed variants of concern (VoC) emerged in the UK [5], in South Africa [6] and in Brazil [7]. These variants, B.1.1.7, B.1.351 and P.1, are more infectious than wild type SARS-CoV-2 and feature extensive changes in both the receptor binding domain (RBD) and the N-terminal domain (NTD) of the spike protein. These two domains harbor the vast majority of neutralizing epitopes [8–14] and consequently it has been observed that – especially for B.1.351 – the neutralizing activity of wild type post-infection and post-vaccination sera is reduced [15–18]. In addition, efficacy and effectiveness of vaccines against B.1.351 has been shown to be somewhat reduced, depending on the type of vaccine platform used [19, 20]. For one currently licensed vaccine, the efficacy against B.1.351 was lost [21]. Updated vaccines based on variant spike sequences are currently being tested by vaccine producers and may be licensed in the future if variants emerge that escape vaccine-induced immunity to an even larger degree. However, the process of updating vaccine antigens to match circulating variants is not as straight forward as it seems. Several variants might circulate simultaneously, making it difficult to choose the right antigen for optimal protection. Of course, multivalent vaccines that include more than one variant antigen can be formulated, but this increases complexity and decreases the amount of vaccine doses that can be manufactured. Understanding the antigenic relationship between variants is therefore of high importance.

Here, we vaccinated mice with recombinant spike proteins from the wild type Wuhan-1 strain, B.1.1.7, B.1.351 and P.1 and assessed the resulting cross-neutralization in the sera. Furthermore, we challenged the animals with wild type strain, SARS-CoV-2/human/USA/USA-WA1/2020 (WA1) of SARS-CoV-2 to determine if variant vaccine antigens would still protect from the prototypic virus. Adjuvanted, recombinant spike proteins were chosen as antigen since they reflect vaccines currently in clinical development by Novavax, Sanofi Pasteur and other vaccine manufacturers.

## Results

### Variant spike proteins induce cross-neutralizing antibodies in the mouse model

First, we vaccinated BALB/c mice twice with adjuvanted recombinant spike proteins of wild type SARS-CoV-2 (Wuhan-1), B.1.1.7, B.1.351 and P.1. Three weeks post boost, the animals were bled and the neutralizing activity of their serum was assessed in a well-established microneutralization assay with authentic SARS-CoV-2 [22]. When tested against the respective virus from which the vaccine antigen was derived, all animals mounted strong neutralizing antibody responses (**Figure 1A**), while negative controls showed no neutralizing activity (**Figure 1B**). The negative control group received an irrelevant control protein, influenza virus hemagglutinin. However, there was a trend towards B.1.1.7 vaccinated animals showing higher neutralizing capacity against homologous virus as compared to the other spike antigens. P.1 seemed to induce the lowest neutralizing activity against homologous viruses. These differences were small and only significant for B.1.1.7 versus P.1.

**Figure 1:**
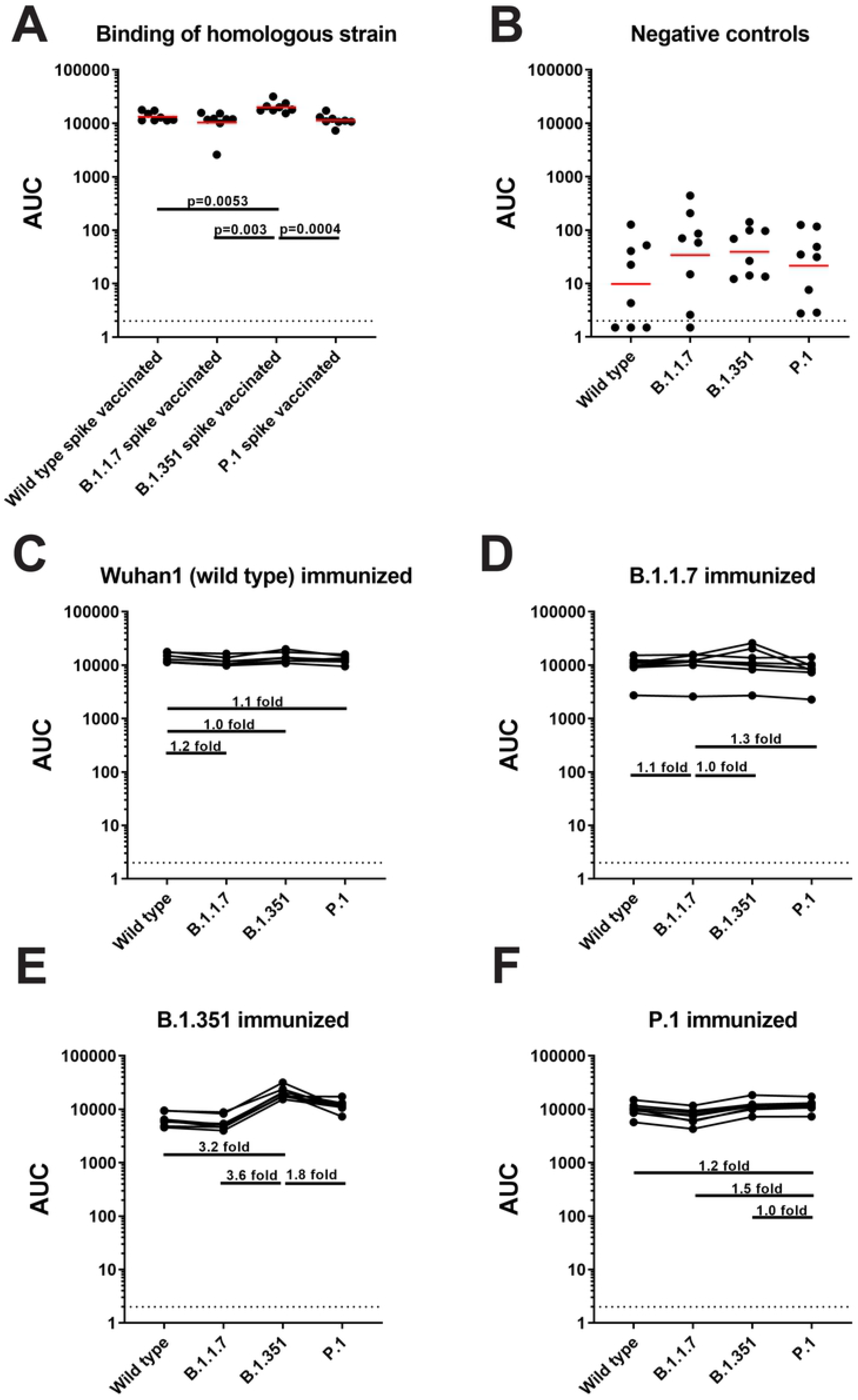
All groups of mice vaccinated with variant spike proteins can cross-neutralize wild type, B.1.1.7, B.1.351 and P.1 isolates of SARS-CoV-2. **(A)** All samples were run on a classic neutralization assay and neutralization activity of each group against homologous strain is shown. **(B)** Serum from the negative control group was also tested against isolates of wild type SARS-CoV-2, B.1.1.7, B.1.351, and P.1 and the ID50s are shown. **(C-F)** Sera from mice vaccinated with wild type spike protein **(C)**, B.1.17 spike protein **(D)**, B.1.351 **(E)**, and P.1 **(F)** was tested against wild type, B.1.1.7, B.1.351 and P.1 isolates of SARS-CoV-2 and the calculated ID50s from neutralization curves are depicted in each graph. The dashed line on each graph indicated the limit of detection (LOD). The differences in neutralization of different variant viruses are indicated by horizontal lines and the fold differences in neutralization are also shown. Statistical significance, when present, is shown as well.

As expected, when testing for cross-reactivity, the different spike proteins induced the highest neutralization titers against the homologous viruses. Sera from wild type spike vaccinated animals neutralized WA1 best, followed by B.1.1.7, P.1 and B.1.351 (**Figure 1C**). Sera from B.1.1.7 vaccinated animals neutralized B.1.1.7 best, followed by wild type, P.1 and B.1.351 (**Figure 1D**). For B.1.351 vaccinated animals, we detected the highest titers against B.1.351 followed by wild type, B.1.1.7 and P.1 (**Figure 1E**). P.1 induced a surprisingly uniform level of immunity with the lowest drop to wild type virus followed by B.1.351 and P.1 (**Figure 1F**). The steepest drops in neutralization were detected for B.1.1.7 to B.1.351 (4.8-fold), from B.1.1.7 to P.1 (4.4-fold), and from B.1.351 to P.1 (4.2 fold). Importantly, we did not observe complete loss in neutralizing activity against any of the viruses.

We used antigenic cartography [23] to visualize the antigenic relationships between the tested viruses and sera (**Figure 2**). The B.1.351 virus is positioned furthest from the WA1 virus, and P.1 and B.1.1.7 are approximately equal distance from WA1 in opposite directions. The sera loosely cluster in the vicinity of the antigen they were raised against.

**Figure 2:**
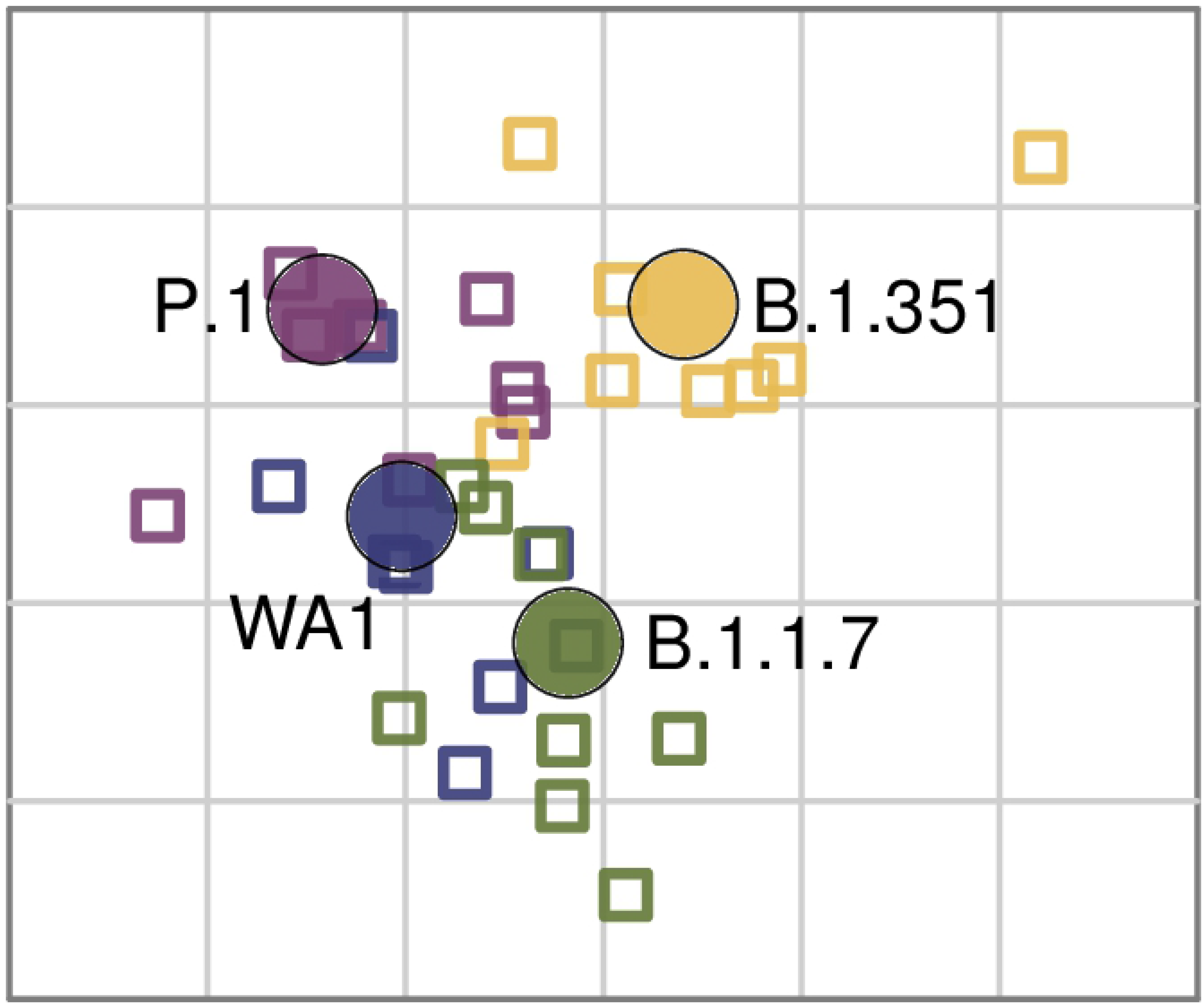
Antigenic map of WA1, B.1.1.7, P.1 and B.1.351 antigens and 31 sera. Antigens are shown as circles (WA1: blue, B.1.1.7: green, P.1: purple, B.1.351: yellow), sera as squares, in the color of the antigen they were raised against. The X and Y axes both correspond to antigenic distance, with one grid line corresponding to a two-fold serum dilution in the neutralization assay. The antigens and sera are arranged on the map such that the distances between them best represent the distances measured in the neutralization assay.

### Antibody binding is less affected than neutralization

We repeated our analysis using an ELISA with the respective spike proteins as substrates. While neutralization requires binding of antibodies to a limited number of epitopes mostly on RBD and NTD, many more binding epitopes exist on the spike protein. Therefore, more even reactivity was expected. We did detect differences in reactivity when binding was tested against the respective matched spikes (**Figure 3A**) but while these differences were statistically significant in three cases, they were relatively small. However, it seemed that vaccination with B.1.351 induced slightly more homologous binding antibodies compared to the other immunogens. Low background reactivity was detected in sera of the control animals (**Figure 3B**).

**Figure 3:**
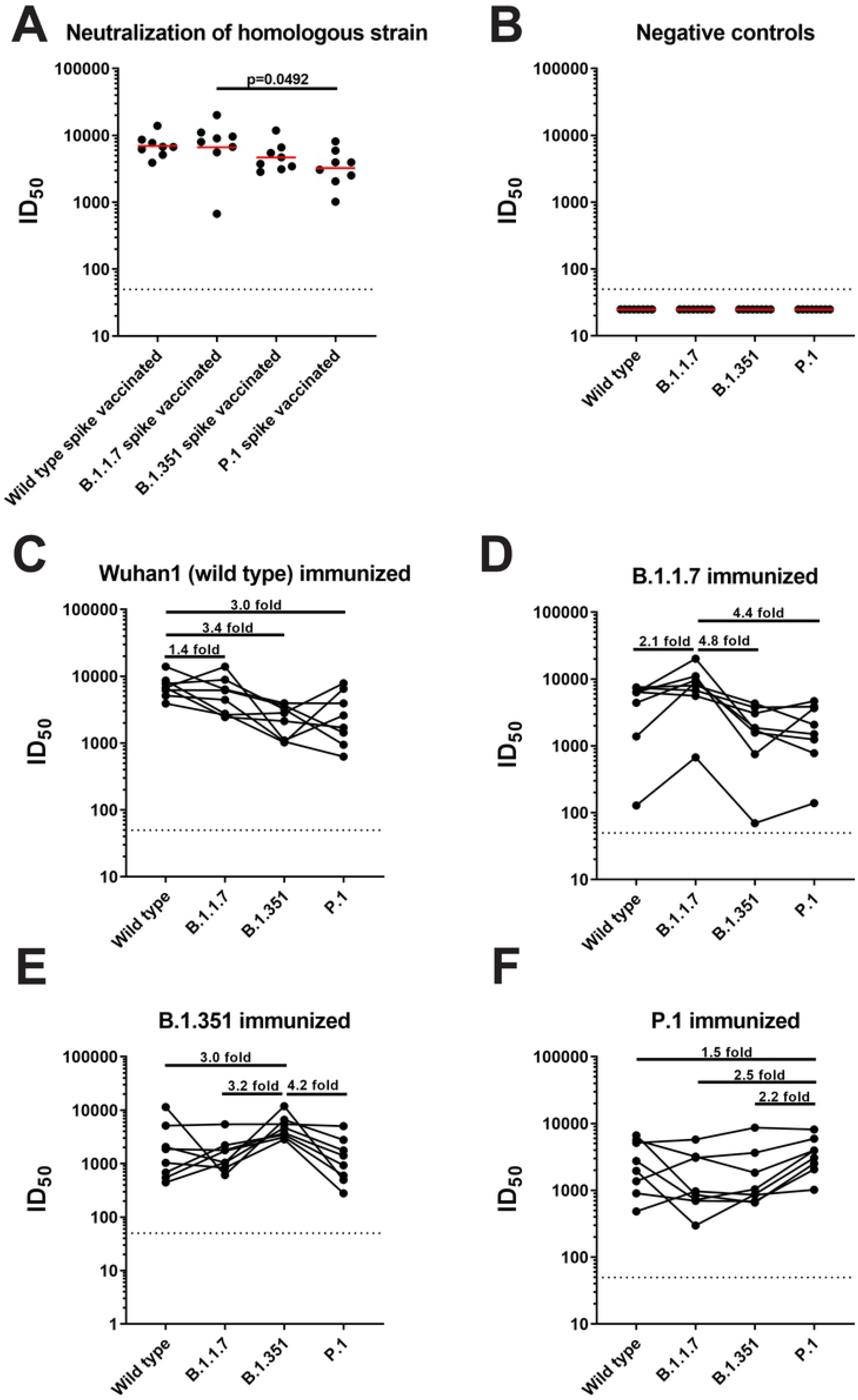
All vaccinated groups have cross-reactive antibodies in their sera against spike proteins of wild type, B.1.1.7, B.1.351 and P.1. **(A)** A standard ELISA was performed using sera from each group and tested for binding with the homologous spike protein and the binding of each group against the respective spike protein is represented as AUC. **(B)** Binding of the samples in the negative control group was also tested against the spike proteins of wild type SARS-CoV-2, B.1.1.7, B.1.351 and P.1 isolates. **(C-F)** Sera from mice vaccinated with wild type spike protein **(C)**, B.1.17 spike protein **(D)**, B.1.351 **(E)**, and P.1 **(F)** was tested against the spike proteins of wild type, B.1.1.7, B.1.351 and P.1. Binding is shown as AUC and the differences in binding are indicated by horizontal bars with the calculated fold increase or decrease. Statistical significance, when found, is shown as well.

Both wild type spike and B.1.1.7 spike induced relatively even binding antibody responses (**Figure 3C and D**) with maximum fold-reduction of 1.2 and 1.3-fold respectively. A stronger reduction was detected when B.1.351 was used as immunogen with 3.2-fold and 3.8-fold reduction in binding to wild type and B.1.1.7 spike respectively (**Figure 3E**). The drop for P.1 was smaller (1.8-fold). P.1 also induced comparable binding antibody response with a maximum fold-reduction of 1.5-fold against B.1.1.7 (**Figure 3F**).

These discrepancies between neutralization and binding antibody profiles allowed us to calculate ratios between binding and neutralizing antibodies. The best (higher) ratios (indicating higher neutralization) were found in sera from wild type and B.1.1.7 vaccinated mice (**Supplementary Figure 1A**). For each vaccination group, the ratio was always best against the homologous virus and dropped with antigenic distance (**Supplementary Figure 1B-E**). The most stable ratio was observed for P.1 vaccinated animals (**Supplementary Figure 1E**).

### All spike vaccinated animals are protected against challenge with wild type SARS-CoV-2

Finally, we wanted to assess if the induced neutralizing antibody responses can protect animals from challenge with prototypic SARS-CoV-2 strain WA1. Since BALB/c mice are not susceptible to this virus, they had to be pre-sensitized via intranasal transduction with adenovirus expressing human angiotensin converting enzyme 2 (hACE2) before challenge, as previously described. The main readout for the challenge experiment were virus titers in the lungs of infected animals. On day 2 post challenge, control animals showed high viral loads in their lungs (approximately 10^6^ plaque forming units per ml of lung homogenate) (**Figure 4A**). In contrast, no virus was detected in wild type and P.1 spike vaccinated animals. For B.1.1.7 and B.1.351 spike vaccinated animals, one animal per group showed traces of virus replication in the lung, but titers were barely above the limit of detection. On day 5 post infection, no virus was detectable in the lungs of vaccinated individuals while control animals still showed high virus loads (**Figure 4B**).

**Figure 4:**
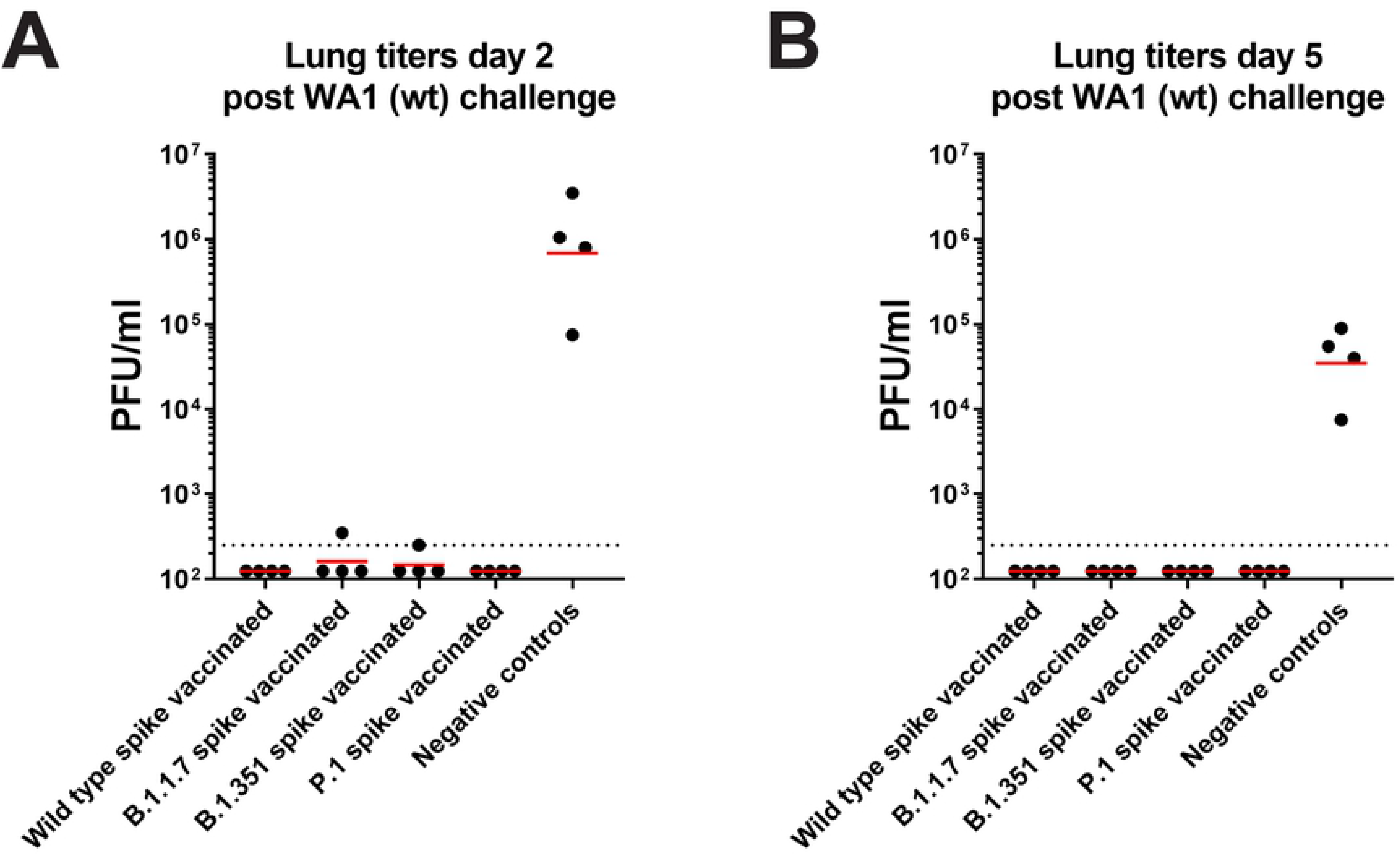
Vaccination with spike proteins of B.1.1.7, B.1.351, and P.1 protects against challenge with wild type SARS-CoV-2 in a mouse model. **(A-B)** All groups of vaccinated mice were challenged with authentic SARS-CoV-2 after sensitization with AdV-hACE2 five days prior to infection. After infection, viral loads in the lungs were quantified via a plaque assay on day 2 **(A)** and day 5 **(B)**.

## Discussion

The emergence of SARS-CoV-2 variants is concerning both in terms of infection control (because many variants are more infectious), as well as in terms of vaccine effectiveness, due to the potential for immune escape. While several vaccines which are either in clinical development, or already in use show good efficacy or effectiveness against currently circulating variants [19, 20, 24, 25], the vaccine antigens may need to be updated at some point if variants emerge that evade neutralizing antibodies more efficiently. However, if variants co-circulate, it will be difficult to select the ‘right’ variant that induces good immune responses across the board. Here, we have tested the cross-neutralization activity of wild type, B.1.1.7, B.1.351 and P.1 adjuvanted protein vaccines in the mouse model. We found that P.1 induces the most balanced immune response across the four tested antigens, supported by four sera raised against P.1 positioned centrally in the map between WA1, B.1.351 and P.1. Interestingly, while B.1.351 and P.1 share two of their RBD mutations and have a mutated residue at position 417 in common (although to different amino acids), a sharp drop in neutralizing activity from B.1.351 to P.1 was observed. However, this relationship was asymmetric, since the drop from P.1 to B.1.351 was much smaller. When considering the results from this study, P.1 should likely be the chosen immunogen for updated vaccines in our mouse model. Our mouse data is similar compared to human cross-neutralization data with the same four variants published by Liu et al. [26]. This suggests that mice may be a good model system to study antigenic variability among variants. Such model systems are of importance, as they ensures the continued availability of first-infection sera for the characterization of novel variants. However, there may be subtle differences between mouse strains and certainly between mice and humans, that need to be further explored. Interestingly, binding was much less affected than neutralization, likely due to many more conserved binding epitopes present on the spike outside of RBD and NTD (which harbor most of the neutralizing epitopes). Importantly, all spike antigens, independently of the lineage, provided robust protection against challenge with the prototypic WA1 strains, suggesting that ‘updated’ vaccines – especially if they induce high neutralizing antibody titers – would sufficiently protect against most other circulating variants as well as prototypic SARS-CoV-2 strains.

Our work here has focused on neutralizing and binding antibodies, which have been implicated as correlates of protection for SARS-CoV-2 vaccine induced immunity [27, 28] and reduction in neutralizing antibodies in sera from convalescent individuals and vaccinees against variants has been observed. However, T-cell responses very likely contribute to protection from COVID-19 as well. We have not analyzed T-cell responses in our experimental animals but others have shown that the impact of variants on these responses is minimal [29]. Another caveat of our study is that we were not able to include B.1.617.2, B.1.617.1 and C.37 in our analysis even though these are currently important variants.

In summary, we found that neutralizing titers are always highest against the homologous virus but that antigenic relationships are not necessarily symmetric and that some variant spike proteins induced more balanced responses (e.g. P.1) than others (B.1.351, B.1.1.7). In addition, the drop in binding antibody is much lower than the drop in neutralizing activity. Non-neutralizing binding antibodies have been shown to play an important role in protection for other diseases caused by virus infections including Ebola virus disease and influenza A and B virus [30–33]. The maintenance of binding antibody and T cell responses against variants could partially explain the maintenance of vaccine efficacy, despite the occasional steep drops in neutralizing antibody titers.

## Materials and Methods

### Recombinant proteins

All recombinant proteins were expressed and purified using Expi293F cells (Life Technologies,) as described in detail previously [34, 35]. The spike gene of each respective variant (EPI_ISL_703454, EPI_ISL_745160) was cloned into the pCAGGS vector and used to transfect cells. The cleavage site was deleted by removing the arginine residues and prolines were added to position 986 and 987 to stabilize the spike trimer. The supernatant was clarified on day 4 post transfection via centrifugation at 4,000 g for 20 mins. Ni-NTA agarose (Qiagen) was used to purify the protein, as described before [36, 37]. EPI_ISL_792680 was cloned into pcDNA3.4 for transient transfection. The endogenous leader peptide was replaced with the tPA secretion signal, 8XHIS and AviTag epitopes were appended, and the substitutions noted above introduced. The spike trimer was expressed by transient transfection in 293F cells and purified by affinity chromatography as previously described (PMID: 33842901).

The proteins used for ELISA were purchased from Sino Biological and include the following: 40589-V08B6, 40589-V08B7, 40589-V08B8 and 40589-V08B1.

### Cells and viruses

Vero.E6 cells (ATCC CRL-1586, clone E6) were kept in culture using Dulbecco’s modified Eagle medium (Gibco) which was supplemented with 10 mL of Antibiotic-Antimycotic (100 U/ml penicillin–100 μg/ml streptomycin–0.25 μg/ml amphotericin B; Gibco), 10% of fetal bovine serum (FBS; Corning), and 1% HEPES (N-2-hydroxyethylpiperazine-N-2-ethane sulfonic acid; Gibco). Wild type SARS-CoV-2 (isolate USA-WA1/2020), hCoV-19/South Africa/KRISP-K005325/2020 (B.1.351, BEI Resources NR-54009), hCoV-19/Japan/TY7-503/2021 (P.1, BEI resources NR-54982) and hCoV-19/England/204820464/2020 (B.1.1.7, BEI Resources NR54000) were cultured in Vero.E6 cells for 3 days at 37°C and then the supernatant was clarified via centrifugation at 1,000 g for 10 mins. Virus stocks were stored at −80°C. The protocol is described in greater detail previously [35, 38]. All work with authentic SARS-CoV-2 was performed in the biosafety level 3 (BSL-3) facility following institutional guidelines.

### *In vivo* mouse studies

All animal procedures were performed by adhering to the Institutional Animal Care and Use Committee (IACUC) guidelines. Six to eight weeks old female, BALB/c mice were vaccinated via the intramuscular route with 3 μg of each respective protein with 1:1 mixture of Addavax (Invivogen) in a total volume of 50 μL. After 3 weeks, mice were bled and vaccinated again. Three weeks later, mice were administered anesthesia via the intraperitoneal route and then intranasally transduced with AdV-hACE2 at 2.5×10^8^ plaque-forming units (PFU) per mouse. Anesthesia was prepared using 0.15 mg/kg of body weight ketamine and 0.03 mg/kg xylazine in water. Five days later, all mice were infected with wild type SARS-CoV-2 intranasally with 1×10^5^ PFU. Mice were humanely sacrificed on day 2 and day 5 for assessment of virus in the lungs. Lungs were homogenized using special tubes and a BeadBlaster 24 (Benchmark) homogenizer [39, 40]. Viral load in the lung was quantified via a classic plaque assay [41].

### ELISA

Ninety-six-well plates (Immulon 4 HBX; ThermoFisher Scientific) were coated with 2 μg/mL of each respective protein with 50 μL/well overnight at 4°C. The next morning, the coating solution was discarded, and each plate was blocked with 100 μL/well of 3% non-fat milk (AmericanBio; catalog no. AB10109-01000) in phosphate buffered saline containing 0.01% Tween (PBS-T). Blocking solution was kept on the plates for 1 hour at room temperature (RT). Serum samples were tested starting at a dilution of 1:50 with 1:5 fold subsequent serial dilutions. Serum samples were added to the plates for 2 hours at RT. Next, the plates were vigorously washed thrice with 200 μL/well of PBS-T. Anti-mouse IgG-HRP (Rockland; catalog no. 610-4302) was used at a dilution of 1:3000 in 1% non-fat milk in PBS-T and 100 μL of this solution was added to each well for 1 hour at RT. The plates were washed thrice with 200 μLwell of PBS-T and dried on paper towels. Developing solution was prepared in sterile water (WFI; Gibco) using SigmaFast OPD (*o*-phenylenediamine dihydrochloride, catalog no. P9187; Sigma-Aldrich), and 100 μL was added to each well for a total of 10 min. To stop the reaction, 50 μL/well of 3 M hydrochloric acid was added and the plates were read in a plate reader, Synergy 4 (BioTek), at an absorbance of 490 nanometers. Data were analyzed in GraphPad Prism 7.

### Neutralization assay

Twenty-thousand Vero.E6 cells were seeded per well in a 96-well cell culture plate (Corning; 3340) 1 day prior to performing the assay. Serum samples were heat-inactivated at 56°C for 1 hour prior to use. Serum dilutions were prepared in 1× minimal essential medium (MEM; Gibco) supplemented with 1% FBS. Each virus was diluted to 10,000 TCID_50_s/mL and 80 μL of virus and 80 μL of serum were incubated together for 1 hour at RT. After the incubation, 120 μL of virus-serum mixture was used to infect cells for 1 hour at 37°C. Next, the virus-serum was removed and 100 μL of each corresponding dilution was added to each well. One hundred μL of 1X MEM was also added to the plates to make a total volume of 200 μL in each well. The cells were incubated at 37°C for 3 days and then fixed with 10% paraformaldehyde (Polysciences) for 24 hours. The next day, cells were stained using a rabbit anti-N antibody (Invitrogen; PA5-81794) as primary and a goat anti-rabbit secondary conjugated to horseradish peroxidase (Invitrogen; 31460). This protocol was adapted from an earlier established protocol [34, 35, 42].

### Antigenic cartography

A target distance from a serum to each virus is derived by calculating the difference between the logarithm (log_2_) reciprocal neutralization titer for that particular virus and the log_2_ reciprocal maximum titer achieved by that serum (against any virus). Thus, the higher the reciprocal titer, the shorter the target distance. As the log_2_ of the reciprocal titer is used, a 2-fold change in titer will equate to a fixed change in target distance whatever the magnitude of the actual titers. Antigenic cartography [23] (Smith et al 2004) was then used to optimize the positions of the viruses and sera relative to each other on a map, minimizing the sum-squared error between map distance and target distance. Each virus is therefore positioned by multiple sera, and the sera themselves are also positioned only by their distances to the viruses. Hence, sera with different neutralization profiles to the virus panel are in separate locations on the map but contribute equally to positioning of the viruses. The antigenic cartography software used was written by Sam Wilks and as free and open-source software from https://www.antigenic-cartoraphy.org.

## Acknowledgments

We would like to thank Dr. Randy A. Albrecht for oversight of the conventional BSL3 biocontainment facility, which makes our work with live SARS-CoV-2 possible. We are also grateful for Mount Sinai’s leadership during the COVID-19 pandemic. We want to especially thank Drs. Peter Palese, Carlos Cordon-Cardo, Dennis Charney, David Reich and Kenneth Davis for their support. We thank Sam Wilks for the antigenic cartography toolkit, and Sam Wilks, Eric LeGresley, and Sina Tureli for deep background on SARS-CoV-2 antigenic variation. This work was partially funded by the Centers of Excellence for Influenza Research and Surveillance (CEIRS, contract # HHSN272201400008C), and by the generous support of the JPB Foundation and the Open Philanthropy Project (research grant 2020-215611 (5384); and by anonymous donors.

---

**Supplementary Figure 1: Neutralization over binding ratio varies for each group. (A)** Neutralization over binding ratio was calculated and depicted for each group against the homologous virus and homologous spike protein. Statistical analysis was performed, and the p values are indicated when statistical significance was present. **(B-E)** Neutralization over binding ratios are shown for groups vaccinated with wild type spike protein **(B)**, B.1.17 spike protein **(C)**, B.1.351 spike protein **(D)**, and P.1 spike protein **(E)**.

